## Supplemental for "Savory Signaling: T1R umami receptor modulates endoplasmic reticulum calcium store content and release dynamics in airway epithelial cells"

| Antibodies Used | Source | Catalogue Number |
|---|---|---|
| anti-T1R1 | ThermoFisher Scientific | PA528771 |
| anti-T1R3 | Abcam | ab150525 |
| **Cell Lines Used** | **Source** | **Catalogue Number** |
| Beas-2B | ATCC | CRL-9609 |
| Primary human bronchial epithelial cells | Lonza | CC-2450 |
| Primary nasal epithelial cells | This Study | N/A |
| **Chemicals Used** | **Source** | **Catalogue Number** |
| Bovine Serum Albumin | Millipore Sigma | A2153 |
| CellEvent Caspase 3/7 Reagent | ThermoFisher Scientific | C10423 |
| DAF-FM diacetate | ThermoFisher Scientific | D23844 |
| Denatonium Benzoate | Millipore Sigma | D5765 |
| Fluo-8 AM | Abcam | ab142773 |
| Forskolin | Millipore Sigma | F3917 |
| Fura-2 AM | ThermoFisher Scientific | F1221 |
| Isoproterenol | Millipore Sigma | I6504 |
| Lipofectamine 3000 | ThermoFisher Scientific | L3000075 |
| MEM Amino Acids | ThermoFisher Scientific | 11130051 |
| Non-Essential Amino Acids | ThermoFisher Scientific | 11140050 |
| Thapsigargin | Cayman Chemical | 10522 |
| UTP | ThermoFisher Scientific | AAJ23160-03 |
| **Recombinant DNA - Function** | **Source** | **Catalogue Number** |
| AKAR4 - intracellular PKA activity | Addgene | 61619 |
| AKAR4-nls - nuclear PKA activity | Addgene | 138217 |
| Flamindo2 - intracellular cAMP | Addgene | 73938 |
| nls-Flamindo2 - nuclear cAMP | Addgene | 73939 |
| **RNAi used** | **Source** | **Catalogue Number** |
| TAS1R1 DsiRNA Kit | Integrated DNA Technologies | hs.Ri.TAS1R1.13 |
| TAS1R3 DsiRNA Kit | Integrated DNA Technologies | hs.Ri.TAS1R3.13 |
| **Taqman Probes for qPCR** | **Source** | **Catalogue Number** |
| TAS1R1 | ThermoFisher Scientific | Hs01547926_g1 |
| TAS1R2 | ThermoFisher Scientific | Hs00541095_m1 |
| TAS1R3 | ThermoFisher Scientific | Hs00877446_g1 |
| UBC | ThermoFisher Scientific | Hs01871556_s1 |

**Table S1.** A list of reagents used in this study.

**a**

### MEM Amino Acids, Essential Amino Acids

ThermoFisher Product Number 11130051

| Amino Acid | 1x Concentration (mM) |
|---|---|
| L-Arginine hydrochloride | 0.60 |
| L-Cystine | 0.10 |
| L-Histidine hydrochloride-H2O | 0.20 |
| L-Isoleucine | 0.40 |
| L-Leucine | 0.40 |
| L-Lysine hydrochloride | 0.40 |
| L-Methionine | 0.10 |
| L-Phenylalanine | 0.20 |
| L-Threonine | 0.40 |
| L-Tryptophan | 0.05 |
| L-Tyrosine | 0.20 |
| L-Valine | 0.40 |

**b**

### MEM Non-Essential Amino Acids

ThermoFisher Product Number 11140050

| Amino Acid | 1x Concentration (mM) |
|---|---|
| Glycine | 0.1 |
| L-Alanine | 0.1 |
| L-Asparagine | 0.1 |
| L-Aspartic acid | 0.1 |
| L-Glutamic Acid | 0.1 |
| L-Proline | 0.1 |
| L-Serine | 0.1 |

**Table S2.** Formulations of 1x MEM AA (a) and 1x NEAA (b) used in this study.

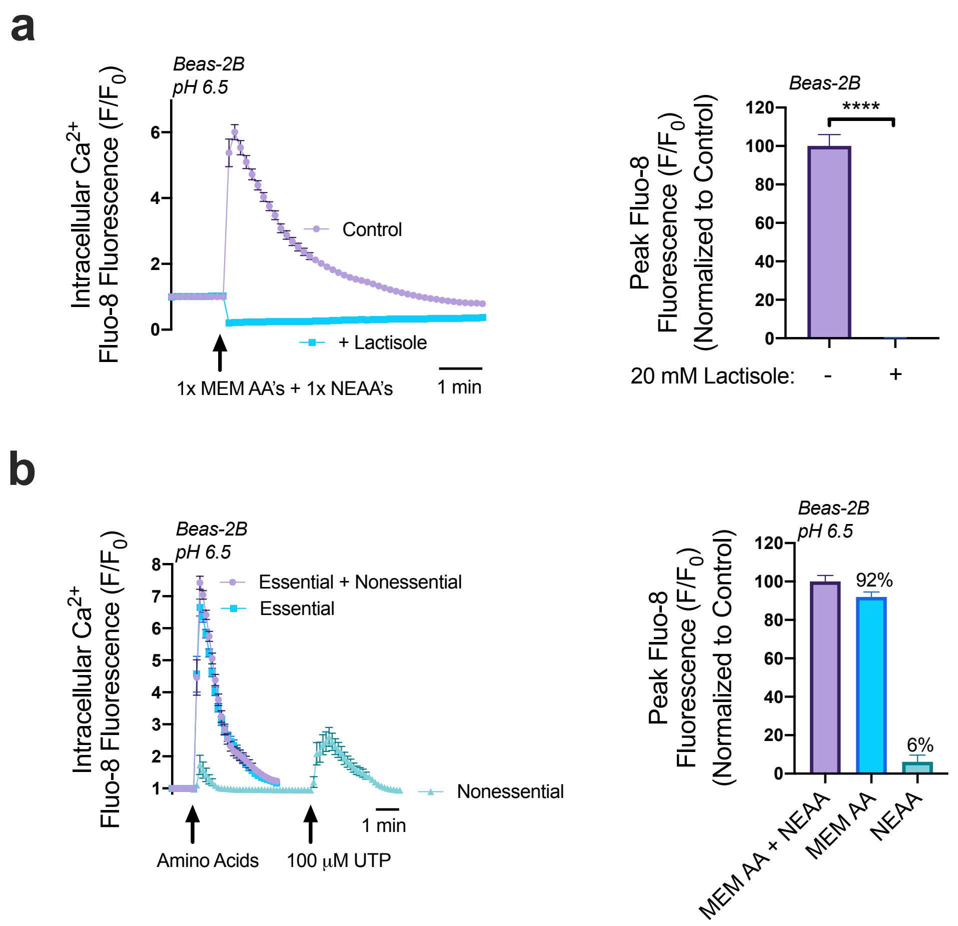


**Figure S1.** Amino acids induce $Ca^{2+}$ elevations in Beas-2B's at pH 6.5. (a) Beas-2B's were loaded with $Ca^{2+}$ detecting dye Fluo-8 AM and pre-treated with 20 mM lactisole for 1 hour then stimulated with a mixture of 1x MEM AA and 1x NEAA's. At pH 6.5 1x MEM AA's and 1x NEAA stimulate a $Ca^{2+}$ response that is inhibited by lactisole. (b) Maintaining a pH of 6.5, the majority of the $Ca^{2+}$ elevations (approximately 92%) are due to components of the 1x MEM AA mixture while 1x NEAA contributed minimally to $Ca^{2+}$ signaling pathways (approximately 6% of the combined release). Significance determined by t-test ****$P < 0.0001$

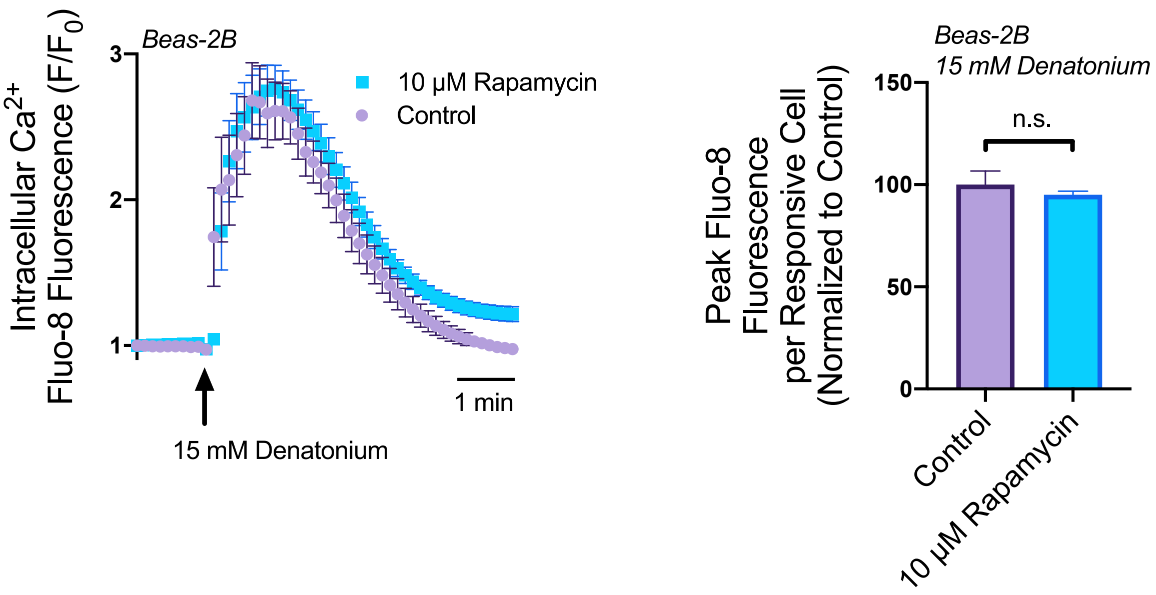


**Figure S2.** mTORC1 pathways do not impact ER $Ca^{2+}$ content. Beas-2B's loaded with Fluo-8 AM were treated with 10 µM of rapamycin for 1 hour then stimulated with 15 mM denatonium. Rapamycin had no impact on the population's peak $Ca^{2+}$ levels or the peak $Ca^{2+}$ release per cell, showing that there was no observable effect on $Ca^{2+}$ signaling pathways. Significance determined by t-test 'n.s.' represents no significance.
